# In Silico Genetics Revealing Novel Mutations in *CEBPA* Gene Associated with Acute Myeloid Leukemia

**DOI:** 10.1101/608943

**Authors:** Mujahed I. Mustafa, Zainab O. Mohammed, Naseem S. Murshed, Nafisa M. Elfadol, Abdelrahman H. Abdelmoneim, Mohamed A. Hassan

## Abstract

**Background:** Myelodysplastic syndrome/Acute myeloid leukemia (MDS/AML) is a highly heterogeneous malignant disease; affects children and adults of all ages. AML is one of the main causes of death in children with cancer. However, It is the most common acute leukemia in adults, with a frequency of over 20 000 cases per year in the United States of America alone.

**Methods:** The SNPs were retrieved from the dbSNP database. this SNPs were submitted into various functional analysis tools that done by SIFT, PolyPhen-2, PROVEAN, SNAP2, SNPs&GO, PhD-SNP and PANTHER, while structural analysis were done by I-mutant3 and MUPro. The most damaging SNPs were selected for further analysis by Mutation3D, Project hope, ConSurf and BioEdit softwares.

**Results:** A total of five novel nsSNPs out of 248 missense mutations were predicted to be responsible for the structural and functional variations of CEBPA protein.

**Conclusion:** In this study the impact of functional SNPs in the CEBPA gene was investigated through different computational methods, which determined that (R339W, R288P, N292S N292T and D63N) are novel SNPs have a potential functional effect and can thus be used as diagnostic markers and may facilitate in genetic studies with a special consideration of the large heterogeneity of AML among the different populations.

## 1. Introduction

Myelodysplastic syndrome/Acute myeloid leukemia (MDS/AML) is a highly heterogeneous malignant disease; affects children and adults of all ages. AML is one of the main causes of death in children with cancer.(1–5) However, It is the most common acute leukemia in adults,(6–8) with a frequency of over 20 000 cases per year in the United States of America alone.(6) It characterized by familial platelet disorder with propensity to myeloid malignancy.(2) AML is frequently caused by mutation in *CEBPα* gene.(9–13) The CCAAT/enhancer-binding protein-alpha (*CEBPα*) is a transcription element that influences immune cell fate and differentiation.(14, 15) Most AML patients with *CEBPα* mutations simultaneously carry double mutations.(16–18) nevertheless, different mutations have been documented.(19–23)Some studies has been reported which claimed some related factors besides *CEBPα* mutation, such as Smoking, alcohol and exposure to solvents and agrochemicals may causes AML, (24–26) but no evidence of publication bias. Other genes have been reported which cause AML such as *FLT3-ITD* and *NMP1* which helped refine individual prognosis. Furthermore, these mutant molecules characterize as a potential targets for molecular therapies.(6, 27–30) Sometimes, chronic lymphocytic leukemia patients can develop AML.(31) While in rare cases AMLs can develop esophageal cancer.(32)

Stem-cell transplantation treatment is associated with the outcome of treatment for patients with cytogenetically normal AML.(33) Nevertheless, the benefit of the transplant was limited to the subgroup of patients with the prognostically adverse genotype consisting of wild-type *CEBPA* without FLT3-ITD.(30, 33) In spite of this hopeful recent evolution, the outcomes of patients with AML remain insufficient, with more than fifty percent of patients eventually dying from this devastating disease. The aim of this study is to identify functional SNPs located in coding region of *CEBPA* gene using in silico analysis.

Disease-causing SNPs are often found to occur at evolutionarily conserved regions. Those have a crucial role for protein structure and function. The ability to predict whether a particular SNP is disease-causing or not is of great importance for the early detection of disease.(34–44) The practice of translational bioinformatics has solid influence on the identification of candidate SNPs and can contribute in pharmacogenomics by identifying high risk SNP mutation contributing to drug response as well as developing novel therapeutic elements for this deadly disease.(45–53) This is the first in silico analysis in the coding region of *CEBPA* gene that prioritized nsSNPs to be used as a diagnostic markers with a special consideration of the large heterogeneity of AML among the different populations.

## 2. Materials and methods

### 2.1. Data mining

The Polymorphic data on SNPs of human *CEBPα* gene was collected from NCBI web site. (https://www.ncbi.nlm.nih.gov/) and the human protein sequence was collected from Uniprot (https://www.uniprot.org/).(54)

### 2.2. Functional analysis

Functional impacts of nsSNPs were predicted using the following in silico tools:

#### 2.2.1. SIFT

SIFT was used to observe the effect of Amino Acids variants on protein on functional level. SIFT predicts damaging SNPs on the basis of the degree of conserved amino A.A. residues in aligned sequences to the closely related sequences, gathered through PSI-BLAST.(55) It is available at (http://sift.jcvi.org/).

#### 2.2.2. PolyPhen-2

PolyPhen-2 stands for polymorphism phenotyping (version 2). PolyPhen-2 was used to observe the influences of Amino Acids substitution on structural and functional level of the protein by considering physical and comparative approaches.(56) It is available at (http://genetics.bwh.harvard.edu/pph2/).

#### 2.2.3. PROVEAN

PROVEAN is a software tool which predicts whether an amino acid substitution has an impact on the biological function of a protein. It is useful for filtering sequence variants to identify nonsynonymous variants.(57) It is available at (https://rostlab.org/services/snap2web/).

#### 2.2.4. SNAP2

SNAP2 is a trained classifier that is based on a machine learning device called "neural network". It distinguishes between effect and neutral variants/non-synonymous SNPs by taking a variety of sequence and variant features into account. It is available at (https://rostlab.org/services/snap2web/).

#### 2.2.5. SNPs&GO

SNPs&GO is an accurate method that, starting from a protein sequence, can predict whether a variation is disease related or not by exploiting the corresponding protein functional annotation. SNPs&GO collects in unique framework information derived from protein sequence, evolutionary information, and function as coded in the Gene Ontology terms, and underperforms other available predictive methods (PHD-SNP and PANTHER).(58) It is available at (http://snps.biofold.org/snps-and-go/snps-and-go.html).

#### 2.2.6. I-Mutant 3.0

I-Mutant 3.0 Is a neural network based tool for the routine analysis of protein stability and alterations by taking into account the single-site mutations. The FASTA sequence of protein retrieved from UniProt is used as an input to predict the mutational effect on protein stability.(59) It is available at (http://gpcr2.biocomp.unibo.it/cgi/predictors/I-Mutant3.0/IMutant3.0.cgi).

#### 2.2.7. MUpro

MUpro is a support vector machine-based tool for the prediction of protein stability changes upon nonsynonymous SNPs. The value of the energy change is predicted, and a confidence score between −1 and 1 for measuring the confidence of the prediction is calculated. A score <0 means the variant decreases the protein stability; conversely, a score >0 means the variant increases the protein stability.(60) It is available at (http://mupro.proteomics.ics.uci.edu/).

### 3. Structural Analysis

#### 3.1. Detection of nsSNPs Location in Protein Structure

Mutation3D is a functional prediction and visualization tool for studying the spatial arrangement of amino acid substitutions on protein models and structures. Further, it presents a systematic analysis of whole genome and whole-exome cancer datasets to demonstrate that mutation3D identifies many known cancer genes as well as previously underexplored target genes.(61) It is available at (http://mutation3d.org).

### 4. Modeling nsSNP locations on protein structure

Project hope is a web-server to search protein 3D structures (if available) by collecting structural information from a series of sources, including calculations on the 3D coordinates of the protein, sequence annotations from the UniProt database, and predictions by DAS services. Protein sequences were submitted to project hope server in order to analyze the structural and conformational variations that have resulted from single amino acid substitution corresponding to single nucleotide substitution. It is available at (http://www.cmbi.ru.nl/hope)

### 5. Conservational analysis

#### 5.1. ConSurf server

ConSurf web server provides evolutionary conservation profiles for proteins of known structure in the PDB. Amino acid sequences similar to each sequence in the PDB were collected and multiply aligned using CSI-BLAST and MAFFT, respectively. The evolutionary conservation of each amino acid position in the alignment was calculated using the Rate 4Site algorithm, implemented in the ConSurf web server.(62) It is available at (http://consurf.tau.ac.il/).

#### 5.2. BioEdit

BioEdit software is intended to supply a single program that can handle most simple sequence and alignment editing and manipulation functions as well as a few basic sequences analyses. It is available for download at (http://www.mbio.ncsu.edu/bioedit/bioedit.html)

## 3. Results

248 missense mutation were retrieved from the dbSNP/NCBI Database, this SNPs were submitted into various functional analysis tools that done by SIFT, PolyPhen-2, PROVEAN and SNAP2. SIFT server represented 28 deleterious SNPs, PolyPhen-2 represented 85 damaging SNPs (29 were possibly damaging and 56 were probably damaging to protein), PROVEAN represented 34 deleterious SNPs while in SNAP2 we filtered the triple-positive deleterious SNPs from the previous three analysis tools, out of 53 There were 19 nsSNPs predicted deleterious SNPs by SNAP2. Table (1) represent the Quarter-positive of deleterious SNPs after filtrations of them the number decreased rapidly to 19 SNP after this submitted them into SNPs&GO and PhD-SNP and PANTHER to run more investigation on this SNP and their effect on the functional level. The triple positive in the three tools were 5 disease-associated SNPs, that shown in Table (2). Finally, we submitted them to I-Mutant 3.0, P-MUT and MUPro respectively (Table 3) to investigate their effect on structural level. All the SNPs were found to cause a decrease in the stability of the protein.

**Table (1):** Affect or damaging nsSNPs associated predicted by various softwares:

| dbSNP rs# | SUB | SIFT<br>PREDICTION | Score | Polyphen<br>prediction | Score | PROVEAN<br>Prediction | Score | SNAP2<br>Prediction | Score |
| --- | --- | --- | --- | --- | --- | --- | --- | --- | --- |
| rs1358286265 | G355V | AFFECT | 0 | Damaging | 1 | Deleterious | -7.083 | effect | 48 |
| rs1402033817 | G355S | AFFECT | 0 | Damaging | 1 | Deleterious | -4.501 | effect | 47 |
| rs1439202716 | R343C | AFFECT | 0 | Damaging | 1 | Deleterious | -6.702 | effect | 43 |
| rs1455027551 | R339W | AFFECT | 0 | Damaging | 1 | Deleterious | -7.798 | effect | 74 |
| rs758726582 | R333C | AFFECT | 0 | Damaging | 1 | Deleterious | -7.409 | effect | 14 |
| rs1422138876 | V328G | AFFECT | 0 | Damaging | 1 | Deleterious | -6.823 | effect | 51 |
| rs1306818311 | R327W | AFFECT | 0 | Damaging | 1 | Deleterious | -6.468 | effect | 47 |
| rs781549846 | R325C | AFFECT | 0 | Damaging | 1 | Deleterious | -6.324 | effect | 19 |
| rs1013241741 | S299C | AFFECT | 0 | Damaging | 1 | Deleterious | -4.866 | effect | 23 |
| rs1391793930 | K298R | AFFECT | 0 | Damaging | 1 | Deleterious | -2.924 | effect | 31 |
| rs1392203731 | K298Q | AFFECT | 0 | Damaging | 1 | Deleterious | -3.899 | effect | 34 |
| rs776590829 | N292T | AFFECT | 0 | Damaging | 1 | Deleterious | -5.788 | effect | 35 |
| - | N292S | AFFECT | 0 | Damaging | 1 | Deleterious | -4.907 | effect | 29 |
| rs1064794962 | R288P | AFFECT | 0 | Damaging | 1 | Deleterious | -6.636 | effect | 66 |
| rs376856647 | E284D | AFFECT | 0 | Damaging | 1 | Deleterious | -2.694 | effect | 44 |
| rs1196766447 | E284A | AFFECT | 0 | Damaging | 1 | Deleterious | -5.588 | effect | 18 |
| rs1352573347 | P233H | AFFECT | 0 | Damaging | 1 | Deleterious | -3.924 | effect | 37 |
| rs1267025311 | R156W | AFFECT | 0 | Damaging | 1 | Deleterious | -2.983 | effect | 71 |
| rs1452063514 | D63N | AFFECT | 0 | Damaging | 1 | Deleterious | -2.992 | effect | 72 |

**Table (2):** Pathogenic nsSNPs associated variations predicted by predicted by various softwares:

| SUB | SNPs&GO |  |  | PANTHER |  |  | PHD-SNP |  |  | P-mut |  |
| --- | --- | --- | --- | --- | --- | --- | --- | --- | --- | --- | --- |
|  | Prediction | RI | Probability | Prediction | RI | Probability | Prediction | RI | Probability | Prediction | Probability |
| R339W | Disease | 1 | 0.539 | Disease | 9 | 0.949 | Disease | 2 | 0.615 | Disease | 0.85 (91%) |
| N292T | Disease | 5 | 0.723 | Disease | 10 | 0.985 | Disease | 4 | 0.749 | Disease | 0.85 (91%) |
| N292S | Disease | 5 | 0.751 | Disease | 10 | 0.98 | Disease | 5 | 0.767 | Disease | 0.85 (91%) |
| R288P | Disease | 4 | 0.704 | Disease | 7 | 0.86 | Disease | 7 | 0.872 | Disease | 0.85 (91%) |
| D63N | Disease | 4 | 0.717 | Disease | 9 | 0.94 | Disease | 4 | 0.697 | Disease | 0.67 (85%) |

**\*RI:** Reliability Index

**Table (3):** stability analysis predicted by I-Mutant version 3.0 and MUPro:

**\*RI:** Reliability Index **Table (3):** stability analysis predicted by I-Mutant version 3.0 and MUPro:
| dbSNP rs# | SUB | SVM2<br>Prediction<br>Effect | RI | DDG Value<br>Prediction | MUpro<br>Prediction | MUpro Score |
| --- | --- | --- | --- | --- | --- | --- |
| rs1455027551 | R339W | Decrease | 3 | -0.21 | Decrease | -0.24675 |
| rs776590829 | N292T | Increase | 2 | -0.35 | Decrease | -1.48124 |
| - | N292S | Decrease | 6 | -0.73 | Decrease | -1.46979 |
| rs1064794962 | R288P | Decrease | 5 | -0.73 | Decrease | -0.66005 |
| rs1452063514 | D63N | Increase | 1 | -0.2 | Decrease | -1.22302 |

## 4. Discussion

This study revealed 5 novel mutations in *CEBPA* gene using different sequence and structure-based algorithms. Figure (1). Single Nucleotide Polymorphisms (SNP) that found in this study could be used in prognostics of disease, because identification of *CEBPA* status in AML has major clinical importance, allowing relapse risk to be stratified properly for post-remission treatment.(63, 64)

**Figure (1):**
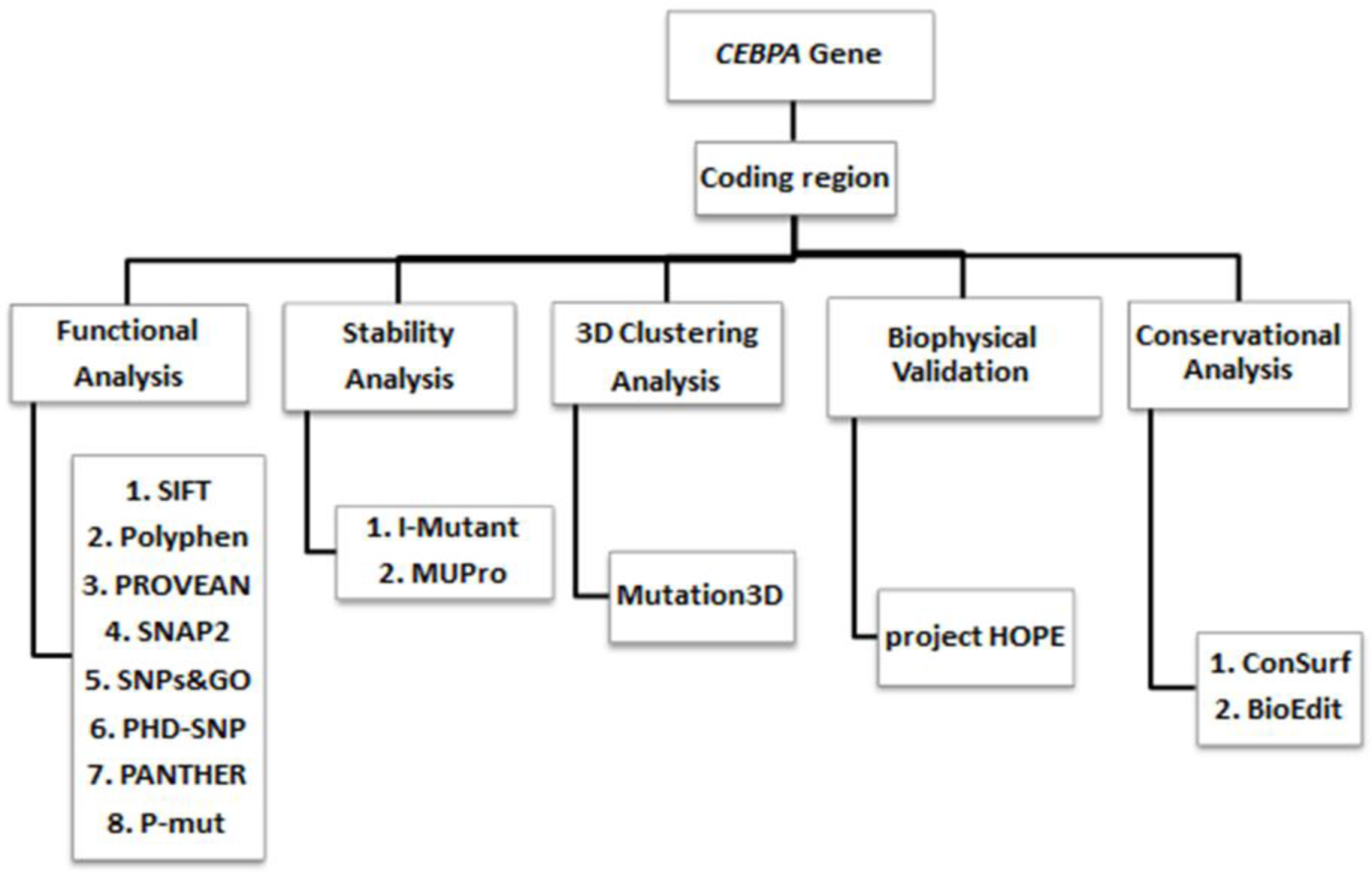
Descriptive Workflow Software’s used in SNPs analysis.

All these SNPs (R339W, R288P, N292S N292T and D63N) were retrieved from dbSNPs/NCBI database as untested, were found to be all novel pathogenic mutations.

*CEBPA* gene provides instructions for making a protein called CCAAT enhancer binding protein alpha. This protein is a transcription factor and it is act as a tumor suppressor, which means that it is involved in cellular mechanisms that help prevent the cells from growing and dividing too rapidly or in an uncontrolled way and that is the principle of cancer (AML). (23, 65)

We also achieved analysis by Mutation3D server, All SNPs in red (R288P, N292S and N292T) are clustered mutation. Significantly, such mutation clusters are frequently associated with human cancers.(66) While SNPs in blue (R339W) and gray (D63N) are covered and uncovered mutations respectively. (Figure 2)

**Figure (2):**
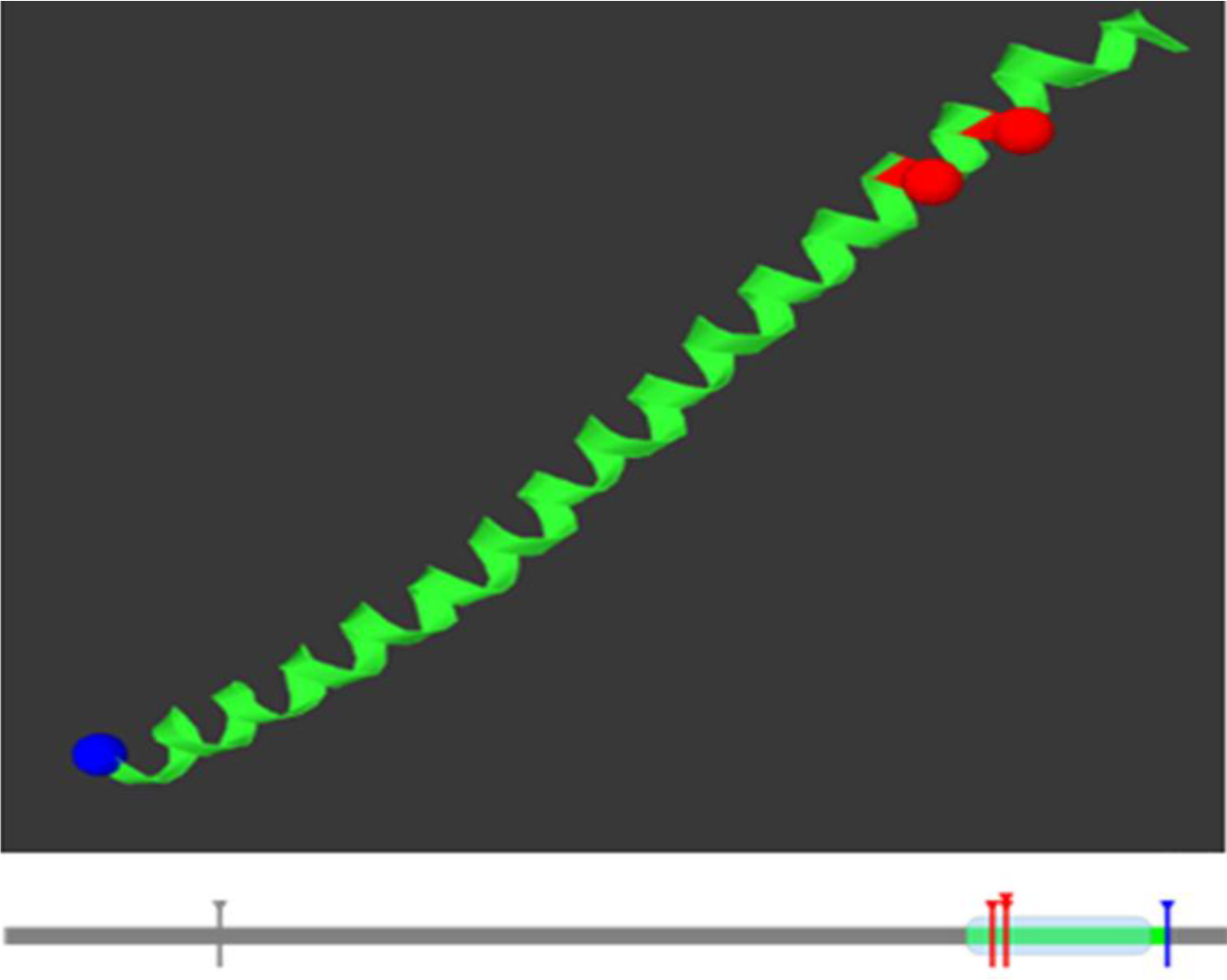
Structural simulations for wild type *CEBPA* gene, demonstrated by Mutation3D.

Project HOPE software was used to submit the most damaging SNPs, (D63N) is the only SNP is located on the surface of a domain, Asparagine (The mutant residue) charge is neutral while Aspartic Acid (the wild-type residue) charge was negative. The charge of the wild-type residue is lost by this mutation; This can cause loss of interactions with other molecules. The mutation is located within a stretch of residues required to repress E2F1:TFDP1-mediated transcription, to inhibit cell cycle and to induce adipocyte differentiation. The differences in amino acid properties can disturb this region and disturb its function. In the second SNP (N292S): The mutated residue is located in a domain that is important for the activity of the protein and in contact with residues in another domain. It is possible that this interaction is important for the correct function of the protein. The mutation can affect this interaction and as such affect protein function. In the third SNP (N292T): The mutation is located within a domain, annotated in UniProt as: bZIP. The mutation introduces an amino acid with different properties, which can disturb this domain and abolish its function. in the fourth SNP (R288P): Proline disrupts an α-helix when not located at one of the first 3 positions of that helix. In case of the mutation at hand, the helix will be disturbed and this can have severe effects on the structure of the protein. (Figure 3)

**Figure (3):**
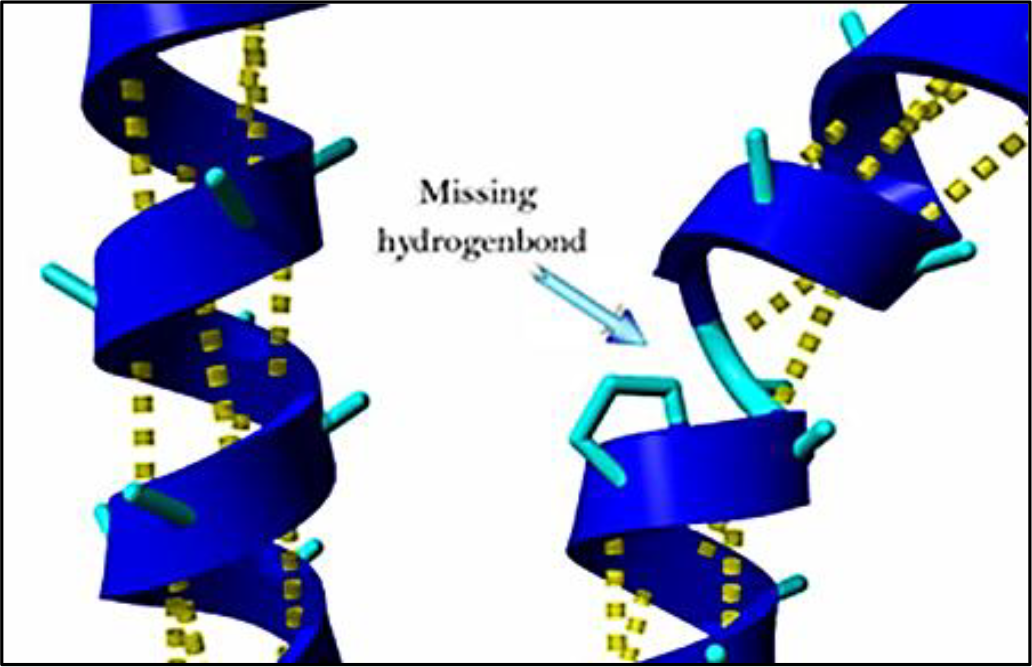
In the 3D-structure can be seen that the wild-type residue is located in an α-helix.

We also used ConSurf web server; the nsSNPs that are showed by black box located at highly conserved amino acid positions have a tendency to be the most damaging nsSNPs. (Figure 4) In this study we also observed that, only one SNP (D63N) the residue predicted to be mutated is evolutionarily conserved across species. (Figure 5)

**Figure (4):**
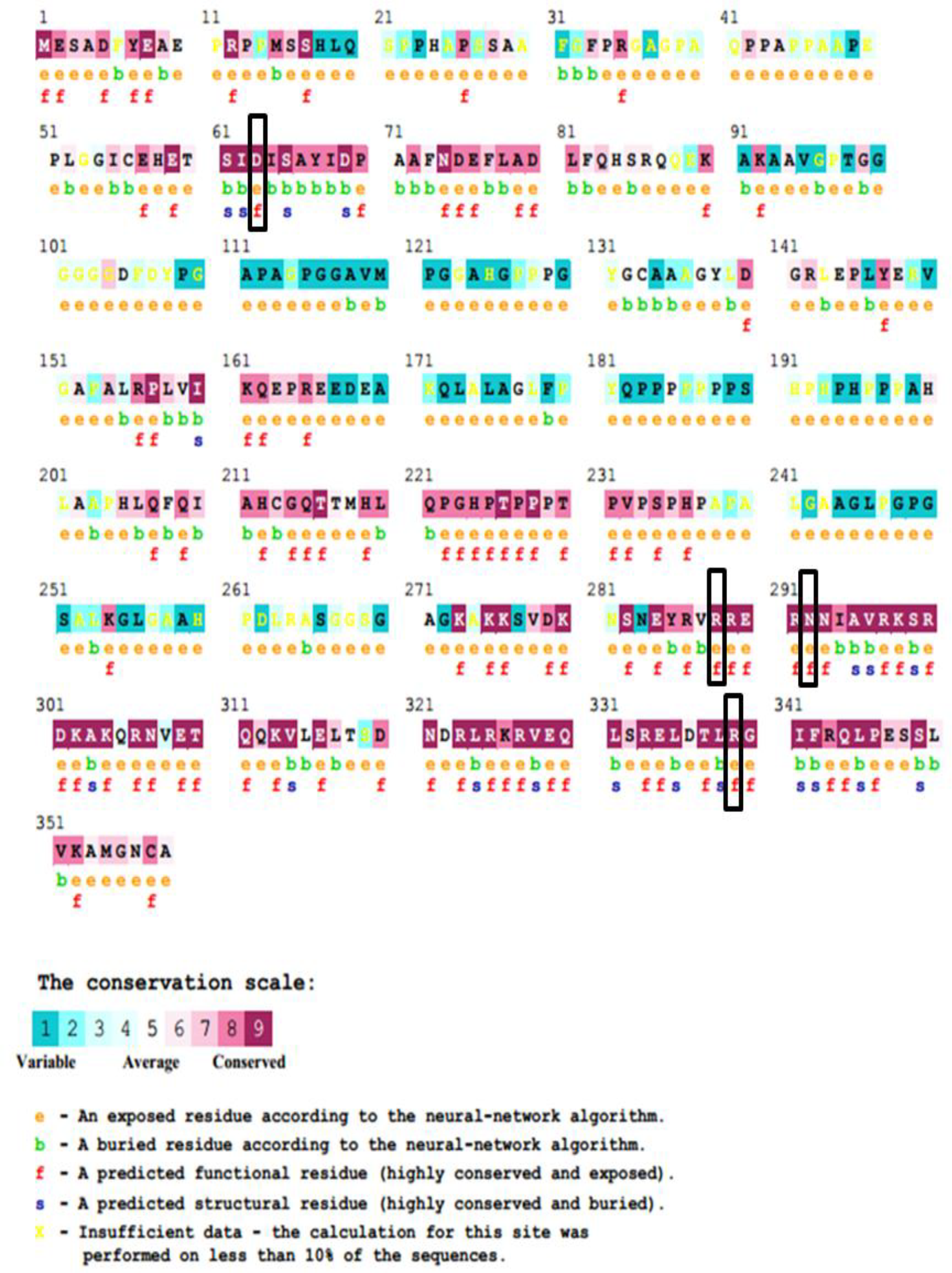
Conserver regions in CEBPA protein were determined using Consurf web server.

**Figure (5):**
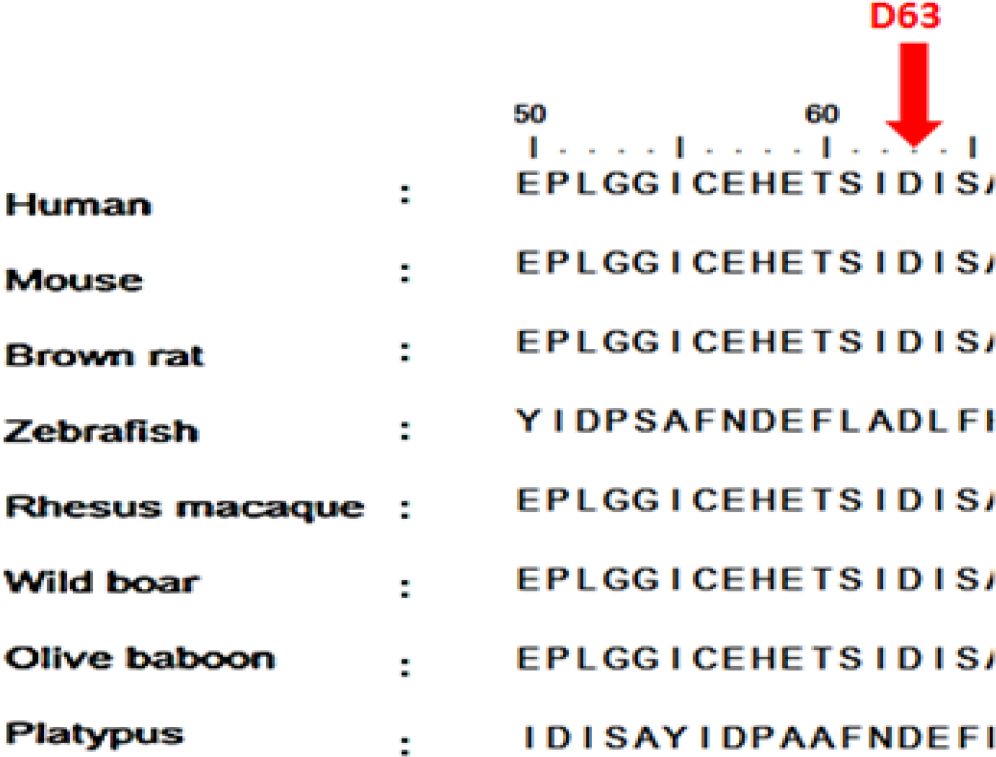
Alignments of 8 amino acid sequences of CEBPA representing that the residues predicted to be mutated are evolutionarily conserved across species.

This study is the first in silico analysis while all other previous studies were NSG analysis, in vitro analysis and in vivo analysis (13, 64, 67, 68)

This study is the first in silico analysis was focused in coding region which revealed five novel mutations that have a possible functional effect and may therefore be used as diagnostic markers for Acute myeloid leukemia and may create an ideal target for cancer therapy. These findings combined with detailed understanding of normal human and murine haematopoiesis makes Acute myeloid leukemia an outstanding model for understanding the principles of cancer progression.(69, 70) Lastly wet lab techniques are recommended to support these findings.

## Conclusion

In this study the impact of functional SNPs in the *CEBPA* gene was investigated through different computational methods, which determined that (R339W, R288P, N292S N292T and D63N) are novel SNPs have a potential functional effect and can thus be used as diagnostic markers. And may facilitate in genetic studies with a special consideration of the large heterogeneity of AML among the different populations. Future studies are needed to confirm those SNPs and identify relevant contributors to disease pathogenesis. This study increased understanding of the pathogenesis of AML by bioinformatics analysis, and may create an ideal target for cancer therapy.

## Data Availability

All data underlying the results are available as part of the article and no additional source data are required.

## Acknowledgment

The authors wish to acknowledgment the enthusiastic cooperation of Africa City of Technology - Sudan.

